# Ultra-high-throughput microbial single-cell whole genome sequencing for genome-resolved metagenomics

**DOI:** 10.1101/2022.04.08.485055

**Authors:** Jie Li, Rong Zhang, Ting Li, Qingyuan Shi, Zhangyue Song, Lichun Jiang, Yunfeng Shi, Conghui Ma, Xue Liang, Jingwei Zhang, Yifan Liu

## Abstract

Microbes exist widely in nature. However, less than 1% of the species of microorganisms have been discovered so far, and more than 99% of microorganisms still remain unknown, which is called microbial dark matter. Unravelling microbial dark matter is not only helpful for exploring the unknown microbial world, but also of great significance for human health research. Ever since a long time ago, due to technical limitations, the microbiome, to a great extent, is still a black box. The traditional population-based metagenomic analysis tools are without single microbial resolution and they are stretched when studying rare and unknown microbial species. Here, we use high-throughput microdroplet technology to achieve the whole genome amplification of microbial single-cell genomes in pico-litter sized microdroplets, and develop subsequent library preparation steps such as indexing of the genome, to achieve the development of ultra-high-throughput microbial single-cell whole genome sequencing for genome-resolved metagenomics, aiming to provide a powerful new weapon for exploring microbial dark matter and other microbiological studies.

## Introduction

According to recent epidemiological and omics-based studies, the microbial communities seem to play a pivotal role in human health and its imbalance in structure and function appears to increase the risk of human disease ^1^. Almost all of microbial communities and humans coexist in different types of symbiotic relationships. A complex and dynamic population of microorganisms resides in gastrointestinal tract and is collectively called gut microbiota, which exerts a remarkable influence on the host during homeostasis and disease^2^. However, due to technical limitations, the gut microbiome has remained a black box to a large extent.

Recently, the popular metagenomic methods^3, 4^, such as 16S rRNA sequencing^5^ and shotgun metagenomic sequencing^6^ have provided comprehensive microbial profiling and putative functions and enabled the exploration of complex microbial communities without need of cultivation. However, these popular metagenomic approaches do not present with a resolution of a single microbe, thus, it is impossible to directly obtain independent data of a single species^7^. The mining of rare and unknown microbial species normally requires obtaining the whole genome information of that species. Thus, a new culture-independent microbiome method with single-cell resolution, especially high-throughput whole-genome sequencing technology, is required to characterize microbial dark matter at the single-cell level.

In recent years, the rapid development of high-throughput single-cell technologies, such as single-cell transcriptome sequencing, single-cell DNA methylome sequencing, and single-cell chromatin accessibility analysis^8-10^, has consequently promoted life science research into the single-cell era. However, such single-cell techniques were mostly applied for the eukaryotic cells^10,11^, and not capable for microbial single cells.

Here, we describe a method to yield high quality genome data from individual microbes through high-throughput sequencing based on droplet microfluidics. Our technique utilizes a hydrogel matrix that enables a series of steps, including washing, lysis of microbes, and genome processing steps. Based on droplet microfluidics, we first lyse the microbes, then perform whole genome amplification, fragment the amplified genomes, and last attach unique barcodes to all fragments. The barcoded fragments of all microbes can then be pooled and sequenced. The sequence reads can be grouped by barcode and a library of single microbe genomes can then be generated, which can be subjected to additional downstream processing, including demographic characterization and in silico cytometry (**Fig.1**).

**Figure 1.**
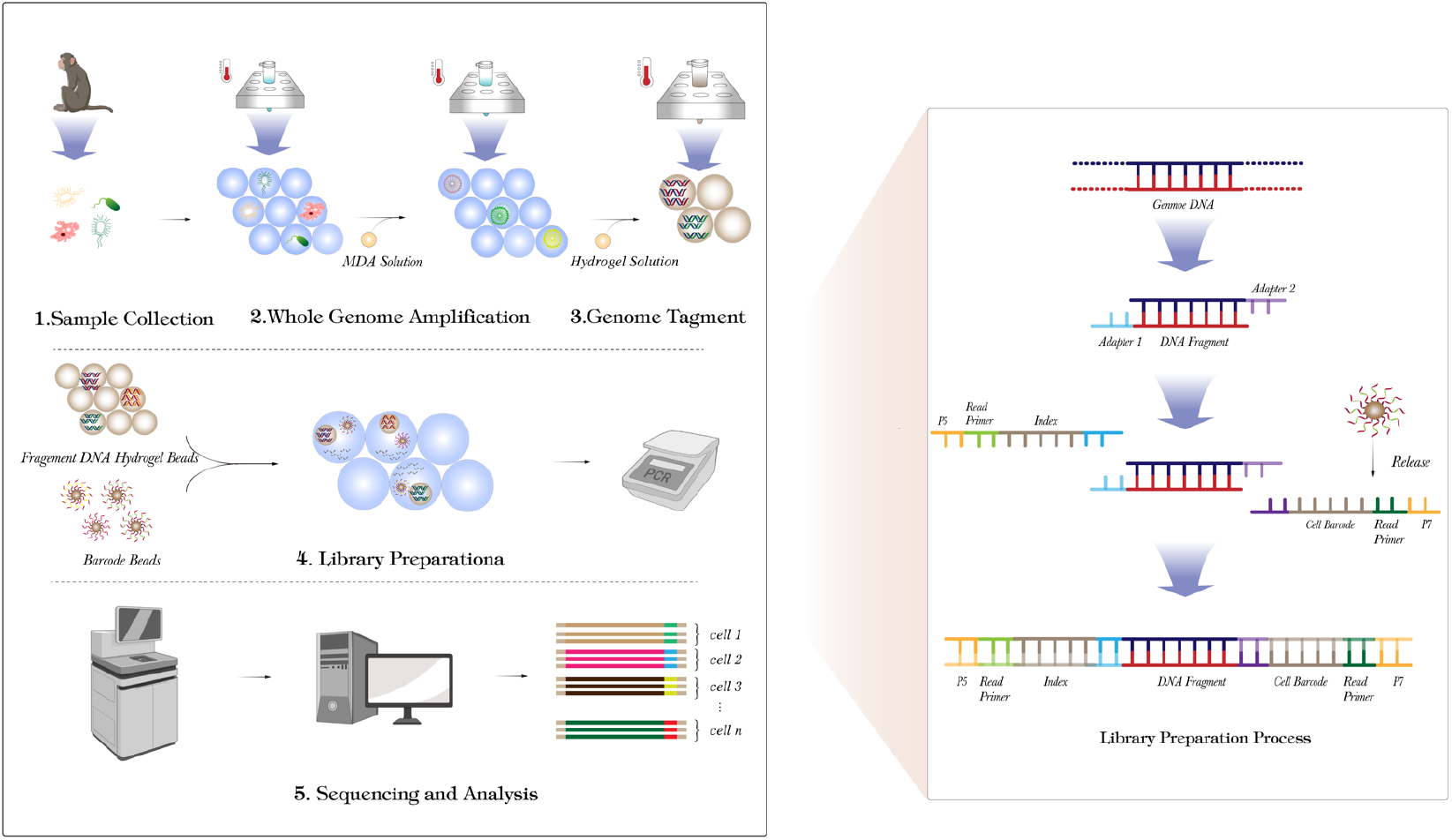
Schematic workflow of microbial single-cell whole genome sequencing.

## Methods

### Microfluidic devices fabrication

The fabrication of the microfluidic device was carried out using polydimethylsiloxane (PDMS)-based soft lithography. First, negative photoresist SU8-3025 (Micro Chem) was coated on a 3-inch silicon wafer (University Wafer) at specific rotating speed to form a uniform and height-specific SU-8 layer as required. The SU-8 treated wafer was then pre-baked at 95°C and the baking time varied depending on the thickness of photoresist. Next, the wafer was exposed to UV lamp (365L2, Thorlabs) to harden the negative photoresist whilst the unexposed areas remained soluble and would be washed away during development in developer solution containing 2-Acetoxy-1-methoxypropane, isopropanol and ethanol (Aladdin). Consequently, the mold was rinsed by isopropanol and ethanol, then dried by nitrogen gun before casting polydimethylsiloxane (PDMS) on it.

A casting approach for creating microfluidic device was performed as following described. A well-mixed 10:1 (base: curing agent) PDMS and precursor (SYLGARD 184, Dow Corning) was casted on the mould followed by baking at 65°C overnight. Then, the cured PDMS chip was cut and peeled off from the mould and punched with a 0.75mm hole puncher to create the inlets and outlets. The fabricated PDMS chip was bonded to a glass slide by plasma bonding, then the chip was baked at 90°C for 2h. Before use, the fabricated microfluidic device was treated with Aquapel silane solution (PPG Industries) to make the channel surfaces hydrophobic.

### Gut microbiota isolation

Gut microbiota was isolated from monkey feces collected by Kunming institute of zoology, Chinese Academy of Science. Fresh fecal samples were instantly frozen at −80 °C without preservatives after sampling. All manipulations as following described were realized under a sterile hood. The frozen fecal samples were first thawed under room temperature under a sterile hood. 5g of monkey feces was diluted and homogenized in 25 ml of normal saline solution (Beyotime) by vertexing at maximum speed for 10 min, followed by a series of filtration steps using membrane filter with minimal pore sizes of 100 μm, 60 μm, 40 μm and 10 μm to remove big aggregates including monkey cells, parasitic ovum and undigested food. The resultant slurry was then centrifuged at 6000 × g for 30 min, and the supernatant was removed carefully. The remaining microbial cell pellets were resuspended in 50% glycerin and stored at −80°C after three times wash in normal saline solution.

### Cell encapsulation and lysis in microdroplets

To prepare the gut microbiota community for processing through the whole workflow, the frozen stock of microbial cells was thawed gently in a room-temperature water bath and washed three times in 1 x PBS buffer (Fisher Scientific, catalog no. AM9624). Microbial cell concentration was determined by manual cell counting under the microscope (Nikon), and diluted in 1 x PBS buffer to an appropriate concentration (10^7^ cells/ml) for single cell encapsulation. For lysing microbial cells in microdroplets, a lysis buffer was prepared by mixing 1M DTT solution (Qiagen, catalog no. 150345) and DLB buffer (Qiagen, catalog no. 150345) in a ratio of 1:11. The microbial single cell suspension and lysis buffer were loaded into a 1 mL syringe and placed on a syringe pump (New Era, catalog no. NE-501) respectively. HFE-7500 fluorinated oil (3M) with 2% (w/w) PEG-PFPE amphiphilic block copolymer surfactant (ThunderBio) was loaded into 5 mL syringe and placed on a syringe pump (New Era, catalog no. NE-501). The cell suspension, lysis buffer, and oil with surfactant were injected into a co-flow droplet maker (**Fig.2a**) at flow rates of 100, 100 and 400 μL/h, respectively, to form microdroplets containing microbial single cell and lysis reagents. Approximately, 100 μL droplets were collected into a 1.5 mL centrifuge tube (Fisher Scientific) and incubated on the heating block (Fisher Scientific) at 65°C for 15min to ensure complete lysis of the microbial single cell.

**Figure 2.**
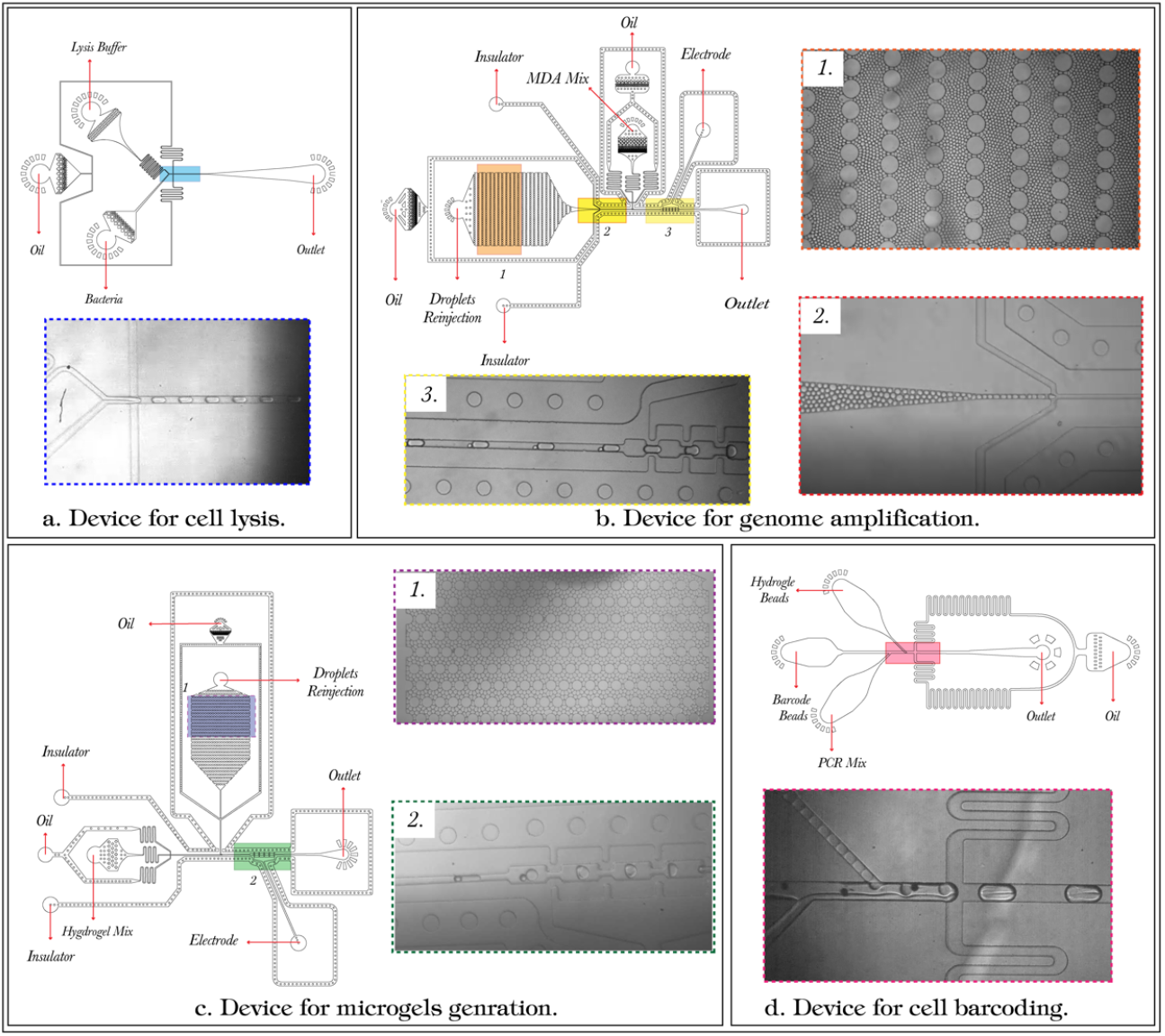
The microfluidics workflow for sequencing library construction. (a) Encapsulation of microbial single cell in microdroplets. (b) Merging the droplets containing microbial single cell genome with MDA reagents in a ratio of 1:1 in order to amplify each microbial single genome in mircodroplets. (c) Merging the droplets containing amplified genomes with precursor hydrogel solution in a ratio of 1:1 for generation of the microgels. (b) Co-encapsulation of microgels containing genome fragments, PCR Mix and barcode beads in microdroplets for indexing all genome fragments of each microbial single cell.

### Whole genome amplification in microdroplets and microgels generation

Multiple displacement amplification (MDA) is used to achieve the whole genome amplification in microdroplets after lysis. 440 μL of MDA reaction mixture containing 290 μL REPLI-g sc Reaction Buffer (Qiagen, catalog no. 150345), 20 μL REPLI-g sc DNA Polymerase (Qiagen, catalog no. 150345), 30 μL 10 x Eva green (biotium, catalog no. 31000), 40 μL Stop Solution (Qiagen, catalog no. 150345), 22 μL N,N-methylenebisacrylamide (Sigma, catalog no.146072) and 38 μL nuclease-free water (Fisher Scientific, catalog no. AM9937) was loaded into 1mL syringe. HFE-7500 oil with 2% (w/w) PEG-PFPE amphiphilic block copolymer surfactant was loaded into 2 mL and 1 mL syringe respectively. The electrode and peripheral insulated channel in the droplets merging devices (**Fig.2b**) were filled with 4M KCL solution. The droplets containing single cell genome after lysis were loaded into a 1mL syringe. The MDA solution, droplets containing cell genome and two syringes of oil were injected into a droplet merging device at flow rates of 200, 25, 200 and 600 μL/h respectively. For generation of droplets containing cell genome and MDA reagents, the electrode syringe was charged with an alternating current (AC) voltage (3 V, 58kHz) to ensure touching droplets containing cell genome and MDA droplets to merge. Approximately, 900 μL of the merged droplets was collected in a 1.5 mL centrifuge tube and then incubated in the heating block at 30°C for 8h to amplify the microbial genomes and 65 °C for 10 min to terminate the amplification reaction. The microdroplets comprising amplified microbial genome and a precursor hydrogel solution containing 40% (v/v) Acrylamide solution (Sigma-Aldrich, catalog no. A4058-100ML) and 20% (w/v) APS (Sigma-Aldrich, catalog no. A9164) were loaded in 1mL syringes respectively and injected with two syringes of oil into a newly designed droplet merging device (**Fig.2c**) at flowrates of 150, 150, 200 and 400 μL/h. Roughly 900 μL droplets were collected into a 1.5 mL centrifuge tube. After adding 0.2 μL TEMED (Sigma-Aldrich, catalog no. T9281-25ML), the collected droplets were kept at room temperature overnight to ensure complete polymerization and solidification of microgels.

### Resuspending microgels in aqueous buffer

To separate emulsion from oil, the oil layer in tube was carefully extracted by a 1 mL syringe and discarded. Emulsions were broken by adding an equal volume of 20% (v/v) 1H,1H,2H,2H-perfluoro-1-octanol (Aladdin, catalog no. H157208-100g) in HFE7500, then mixed by pipetting and centrifuged at 380 × g for 1min.The oil was removed from the centrifuge tube using a syringe. Following droplet breaking, in case of dissolving and removing the remaining oil, 200 μL n-Hexane (Aladdin, catlog no. H140380-1L) with 1% (v/v) span80 (Sigma-Aldrich, catalog no. S6760-250ML) was added to microgels to dissolve any remaining oil. And this solution was mixed by pipetting and centrifuged at 380 × g for 1 min. After removing the remaining oil, the microgels were washed in 500 μL Tris 7.0 buffer (Fisher Scientific, catalog no. AM9850G) containing 0.1% (v/v) Triton X-100 (Anatrace, catalog no. APX100) nonionic surfactant and 500 μL Tris 7.0 buffer containing 0.1% (v/v) Tween-20 (Sigma, catalog no. P1379-25ML) three times respectively. The resultant microgels comprising amplified microbial genomes were resuspended in Tris 7.0 buffer and stored in 4°C.

### Tagmentation of genomic DNA in microgels

Before tagmentation, the microgels were washed three times in Tris 7.0 buffer. Using reagents from a TruePrep ® DNA Library Prep Kit V2 for Illumina (Vazyme, catalog no. TD501), the washed microgels containing amplified microbial genomes were simultaneously fragmented and tagged with a common adapter sequence according to manufacturer’s protocol. 125 μL microgels containing amplified genomes were mixed with 100 μL 5 x TTBL, 100 μL TTE Mix V50 and 225 μL nuclease-free water and then incubated on a heating block at 55°C for two hours to ensure the adequate fragmentation of the amplified genomes in microgels. After tagmentation, the microgels were washed and resuspended in Tris 7.0 buffer, subsequently stored in 4°C.

### Barcode beads synthesis and quality control

Synthesis of barcode sequences on the polystyrene beads was achieved through a split-pool approach. Firstly, poly-T and disulfide bonds were synthesized on the surface of polystyrene beads as crosslinker for barcode oligonucleotides. Subsequently, synthesis of barcode oligonucleotides (5’-CAAGCAGAAGACGGCATACGA-NNNNNNNNNNNN-GTCTCGTGGGCTCGG-3’) which contains 12 bases as cell barcodes was accomplished by a split-pool approach in DNA/RNA synthesizer (Applied Biosystems, catalog no. 3900). In each split-pool round, the polystyrene beads were randomly distributed in four DNA synthesis columns and the beads were labeled with specific barcodes. The resultant barcode beads were removed from the columns and incubated in ammonium hydroxide overnight. To verify the sequence on barcode beads, three probes were designed to capture three different regions in the barcode sequence (5’-FAM-GCCGTCTTCTGCTTGGAAAAAAA-3’; 5’-FAM-TCGTATGCCGTCTTCTGC - 3’; 5’-FAM – CCGAGCCCACGAGAC - 3’). Barcode beads (1000/reaction) were added to a 50 μL of probe hybridization reagent containing 1 x PBS, 0.1ng/mL BSA (Solarbio, catalog no. A8020), 2 μM probe solution and thermal cycled with the following program: 65°C for 5min, 48°C for 8min, 40°C for 8min, 30°C for 8min. After hybridization, the barcode beads were washed three times in 6 x SSC buffer (Sigma, catalog no. S0902-1L) and loaded into Countess (Thermo-Fisher Scientific, catalog no.) for observation under fluorescence microscope (Nikon).

### Microfluidic barcoding of encapsulated microbial cells in microdroplets

Tagmented microgels, 200 μL barcode beads solution containing 20,000 barcode beads, 12 μL N5xx (Vazyme, catalog no. TD202), 6 μL 0.2% SDS (Sigma, catalog no. 74255-250G), 30 μL 10 x PCR stabilizer, 50ul OptiPrep™ (Sigma, catalog no. D1556-250ML), 15 μL 10 x Eva Green, 87 μL nuclease-free water, and 100 μL PCR solution containing 60 μL 5 x TAB (Vazyme, catalog no. TD501), 6 μL TAE (Vazyme, catalog no. TD501), 30 μL 100mM DTT (solarbio, catalog no. D1070-5), 4 μL nuclease-free water were each loaded into a 1 mL syringe and injected into the sequential microfluidic device as shown in Fig.2d. The fluorinated oil (HFE7500) with 5% (w/w) PEG-PFPE surfactant was loaded into a 2 mL syringe as continuous phase of the emulsion. Approximately 500 μL droplets were collected and thermal cycled with the following program: 72°C for 3min, 98°C for 30s, then 15 cycles of: 98°C for 15s, 60°C for 30s, 72°C for 3 min, followed by 72°C for 5 min. After thermal cycling, the emulsion was broken by adding equal volume of 1H,1H,2H,2H-perfluoro-1-octanol, and briefly centrifuged. The supernatant solution was collected and the DNA library was confirmed by Flashgel (Lonza, catalog no. 57023), and then purified using a Zymo DNA Clean& Concentrator™-5 kit (Zymo Research, catalog no. D4014). Using VAHTS DNA Clean Beads (Vazyme, catalog no. N411-02), the DNA library was size-selected for DNA fragments in the range of 300 - 600 bp and quantified by Bioanalyzer 2100 instrument (Agilent) and High Sensitivity DNA chip (Agilent, catalog no. 5067-4626), and sequenced on an Illumina NovaSeq.

### Data analysis

Raw reads from the NovaSeq-generated FASTQ files were filtered by quality and the host contamination was removed by kneaddata. The remaining reads were grouped by barcode sequence using bbtools. A given read was discarded if more than 20% of its bases with a Q-value less than Q20. A given barcode group containing less than 1000 reads were discarded. The resulting reads were classified by taxonomic identities from the Kraken database using Kraken v2.1.2 and the MetaPhlAn database using MetaPhlAn 2.0 to barcode groups by a majority rule. Distribution of barcode groups was calculated using the script *counting*.*py*. Alpha and beta diversities of the gut microbiota were analyzed using Qiime2 v2022.2.

## Supporting information

Results

---

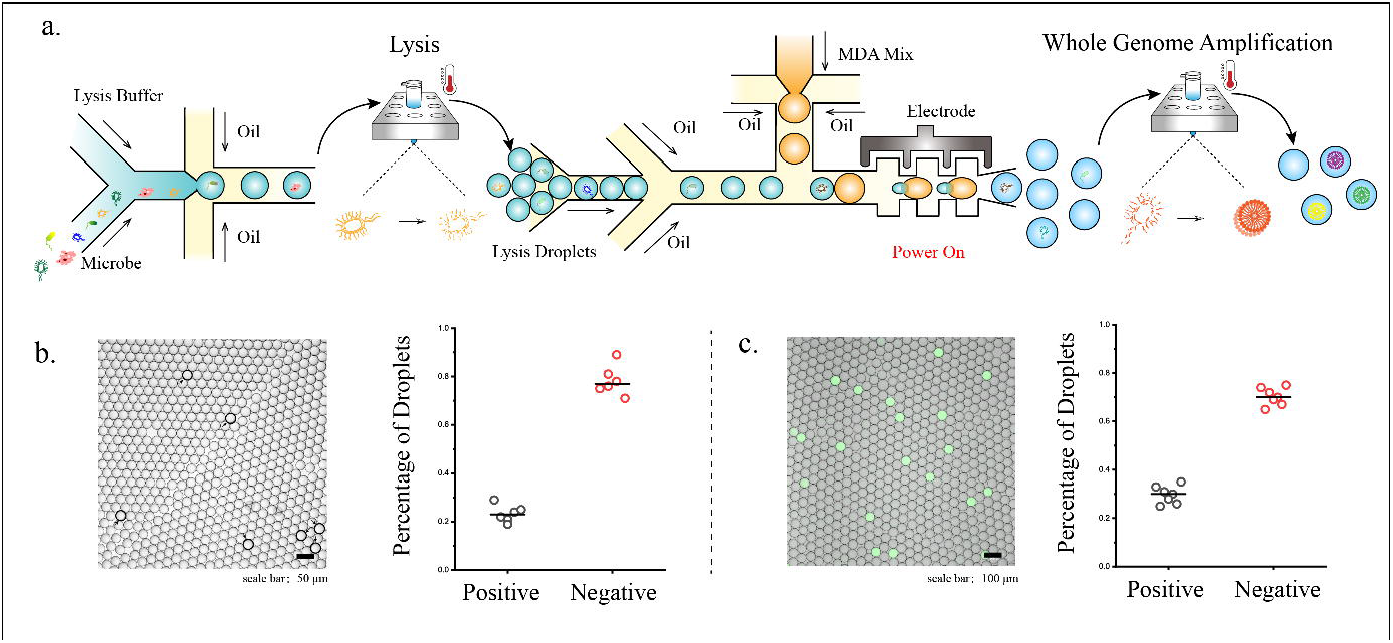

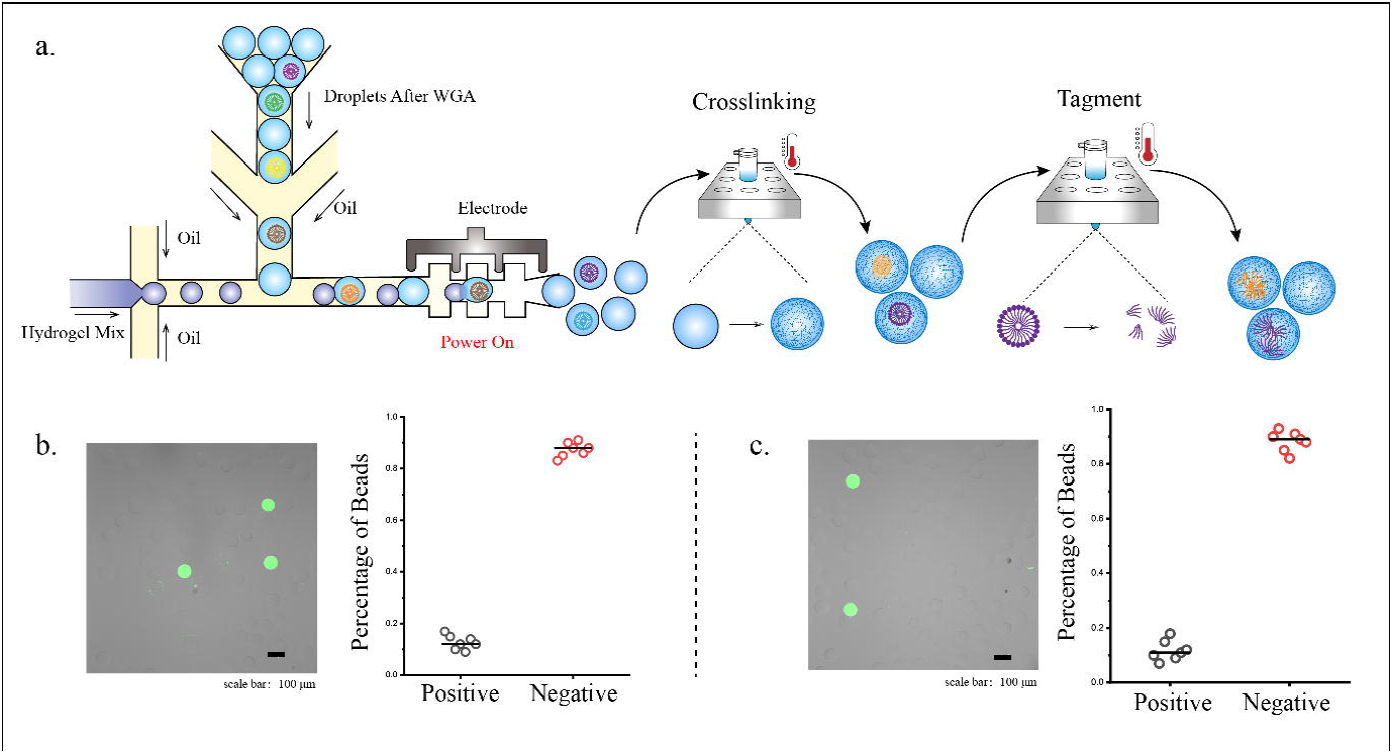

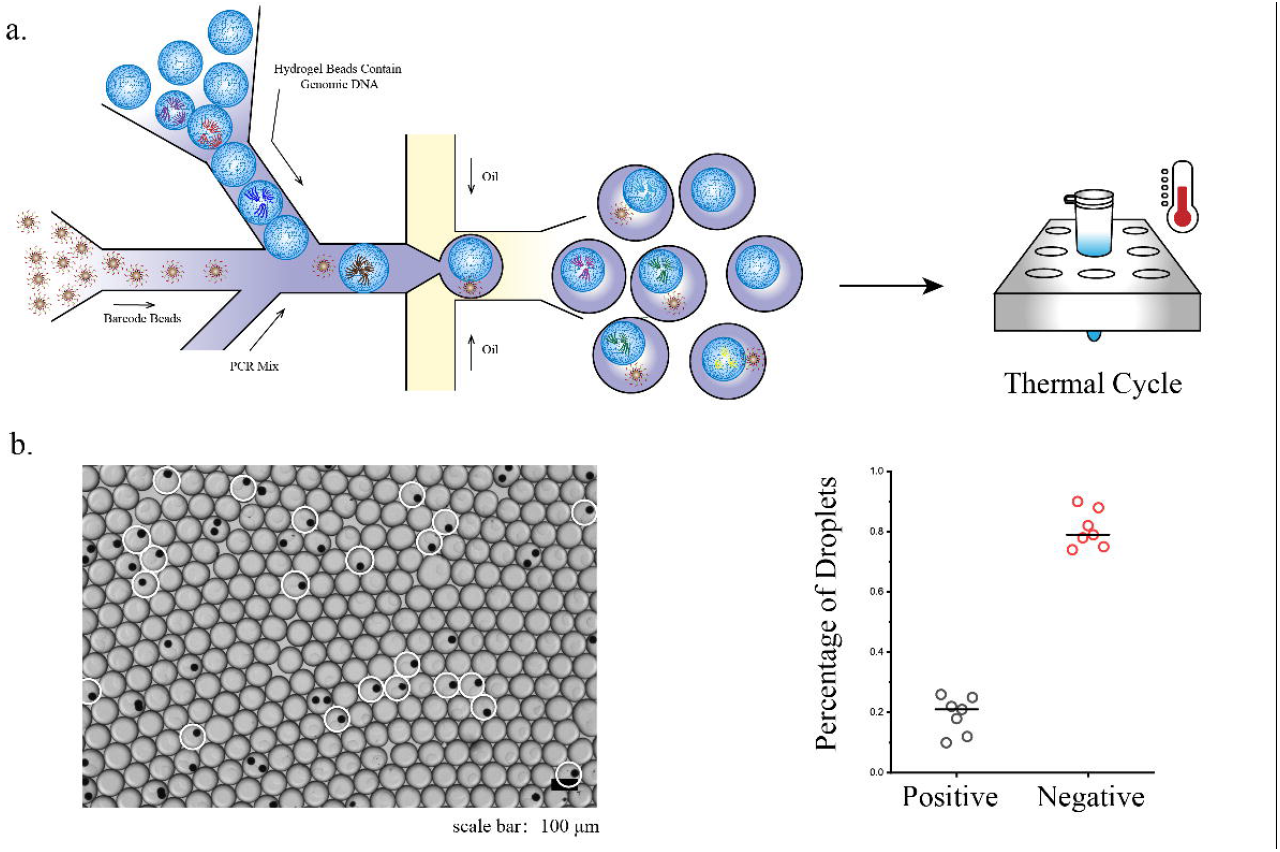

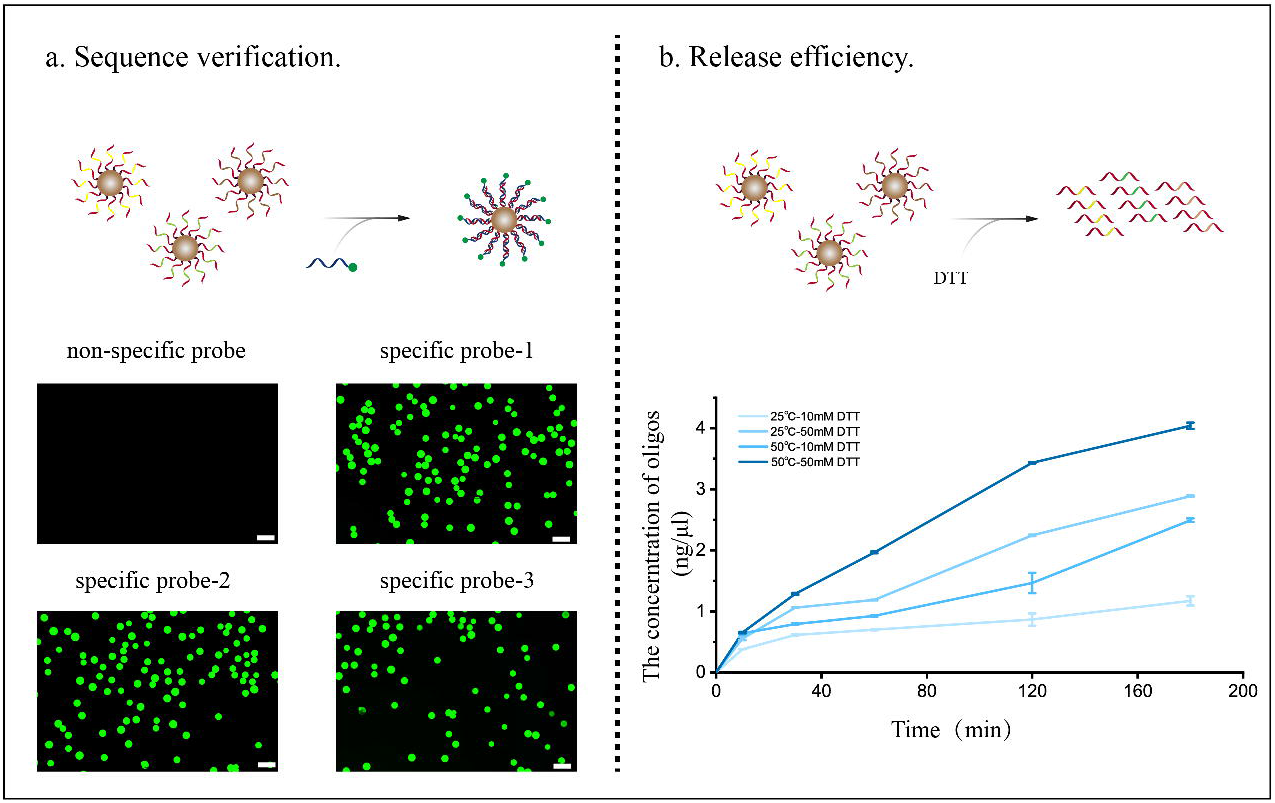

