## Supplementary material for "Ultra-high-throughput microbial single-cell whole genome sequencing for genome-resolved metagenomics": Results

The principal concept of this technique described here is to perform massively parallel microbial single-cell whole genome amplification firstly and then label all DNA fragments originating from the same genome with a barcode sequence which is unique to each microbe so that the resultant products comprise a barcode sequence hybridized to a random genome fragment of each microbe.

To perform massively parallel microbial single-cell whole genome amplification, we first encapsulated each microbial single-cell in lysis buffer-containing droplets using a two-stream co-flow droplet maker to ensure each droplet containing no more than one microbial single-cell. The droplet maker generated about 10,000 droplets per second, allowing us to obtain 30 million 20-μm-diameter droplets in one hour. To lyse microbial single-cell in droplets thoroughly, we incubated the collected droplets at 65℃ for 15min to obtain high molecular weight genomic DNA, which we verified by staining with Eva green dye (**Fig 3b**).

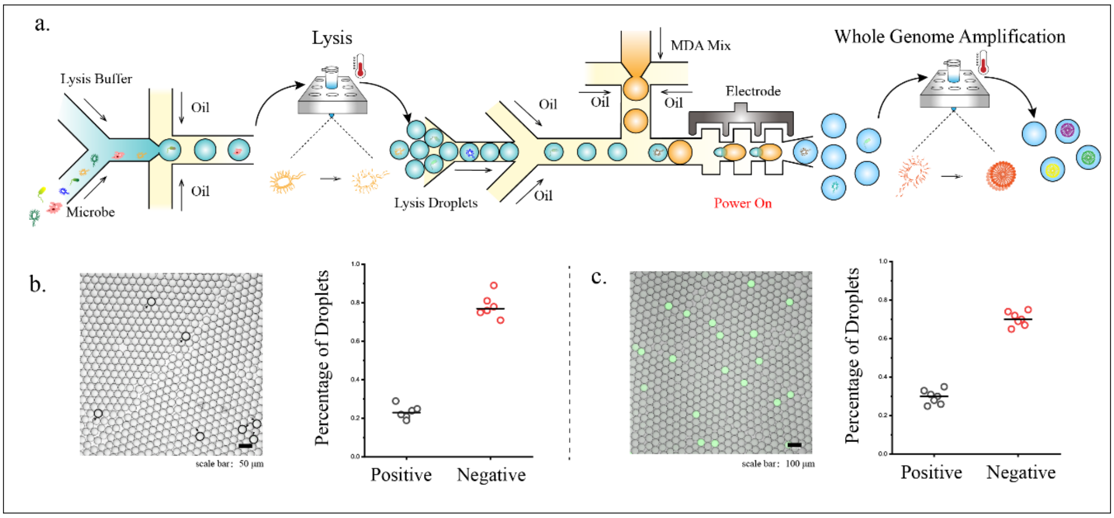


**Fig3: Encapsulation of microbial single-cell, cell lysis and whole genome amplification in droplets.** a. Schematic diagram of encapsulation of microbial single-cell, cell lysis and whole genome amplification in droplets. b. The genomic DNA released after cell lysis was stained with Eva green dye. Positive droplets containing genomic DNA from only one microbial single-cell were highlighted by blank circle; negative means the droplets containing 0 microbial single-cell. c. Whole genome amplification of microbial single-cells was performed in droplets and the amplified genomes were stained with Eva green dye: droplets containing amplified genomic DNA were classified as positive, whereas the others were regarded as negative.

To amplify the microbial single-cell whole genome, we merged the genome-containing droplets with MDA reagents-containing droplets (~ 48μm). The merged droplets were thermal cycled at 30°C for 8h to amplify the whole genome of each microbial single-cell parallelly **(Fig 3c).** After whole genome amplification, the droplets containing amplified genomes were then merged with pre-prepared hydrogel solution-containing droplets in ratio of 1:1 to trap the amplified genomes in solidified hydrogels **(Fig 4b)**, so that the fragmentation of genomic DNA could be simply conducted in bulk without any need for microfluidics. The fragmentation of genomic DNA was succeeded by the “cut and paste” function of Tn5 transposases, which can randomly insert adaptors into DNA and its resulting DNA is ready for downstream reactions. Because of the dimeric nature of Tn5 transposases, the fragmented genomic DNA still remained intact as a macromolecular complex and continued to be retained within the network of solidified hydrogels (Fig 4c).

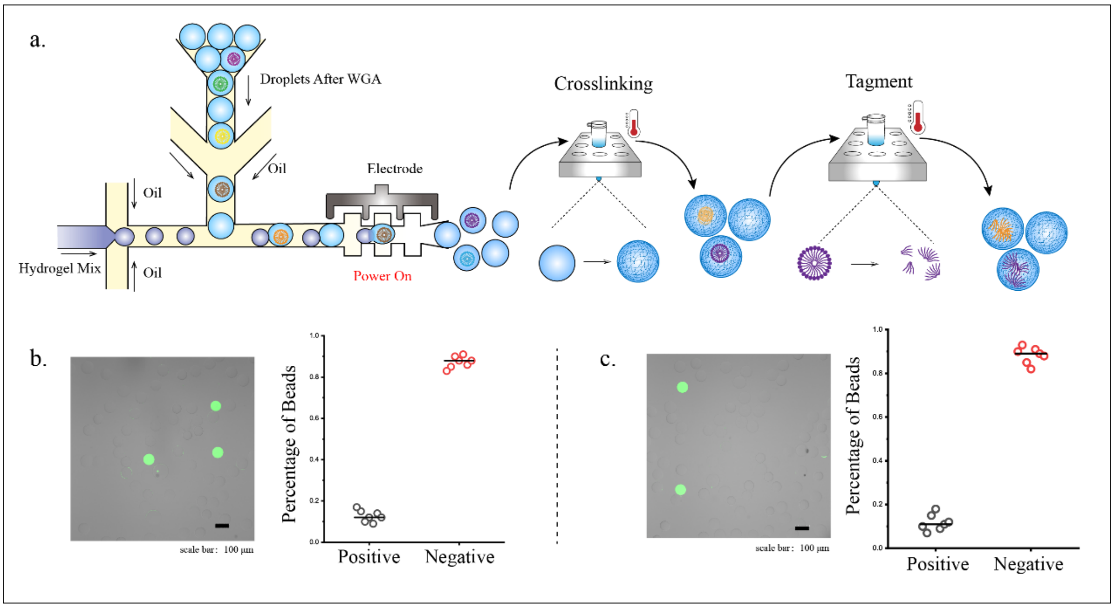


**Figure 4: Trap the amplified genomes of microbial single-cell in solidified hydrogels and fragmentation of genomic DNA with Tn5 transposases in bulk.** a. Schematic plot of generation of solidified hydrogels containing amplified genomes and tagmentation with Tn5 transposases in bulk. b. Positive solidified hydrogels containing amplified genomes exhibited green fluorescence after staining with Eva green dye. c. Positive hydrogels after fragmentation with Tn5 transposases were stained with Eva green dye.

To achieve microbial single-cell whole genome sequencing and distinguish each given sequence read to a given microbial single-cell, we generated libraries of barcode beads, which contained the oligonucleotides covalently binding to the beads. The oligonucleotides comprised 12 random bases flanked by constant sequences with PCR reagents and primers complementary to the constant regions of the barcodes with one side containing the Illumina P7 flow cell adapter. Using droplet microfluidics, we encapsulated DNA fragments-containing hydrogels with barcode beads in PCR reagents and thermal cycled to ensure that the fragmented genomic DNA of each individual microbial single-cell was associated with a unique and identifying barcode sequence (Fig 5). In this instance, all DNA fragments contained both P5 and P7 Illumina sequencing adaptors. The resultant sequencing libraries were then converted into DNA Nanoballs (DNBs) for sequencing with BGI's proprietary DNB'seqTM NGS technology by JMDNA company. After sequencing, the reads were filtered by quality and grouped by each unique and identifying barcode, accommodating us with microbial single-cell whole genome sequence data.

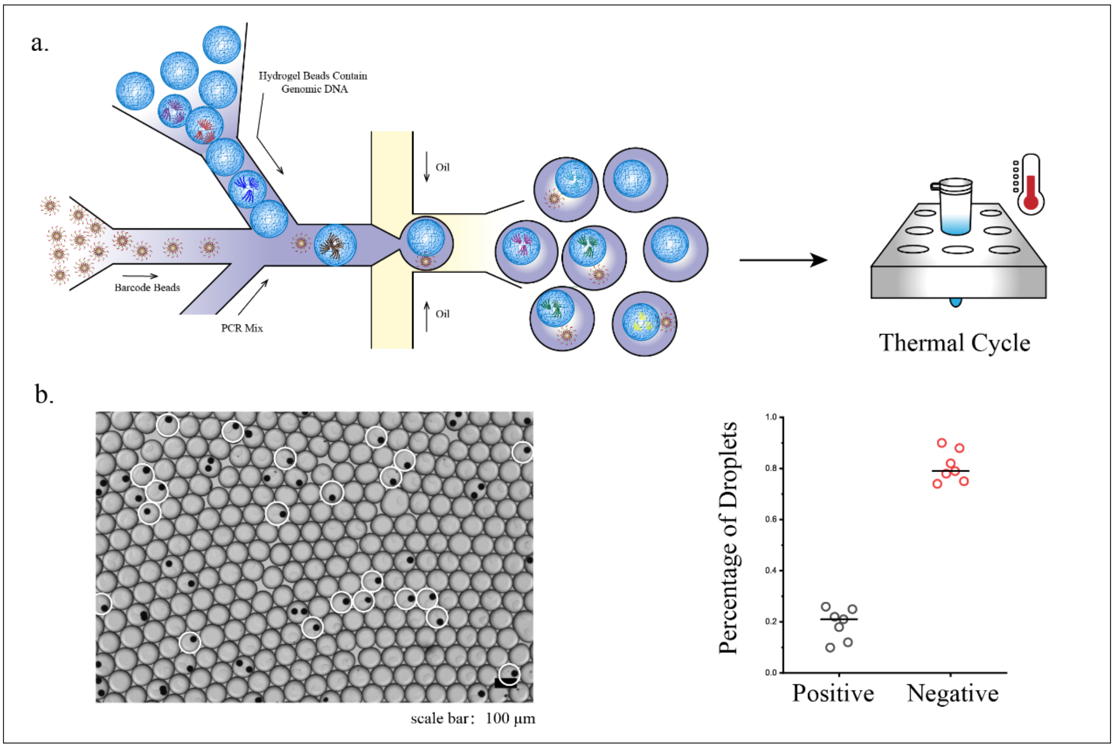


**Figure 5: Co-encapsulation of microbial single-cell hydrogel beads and barcode beads.** a. Schematic diagram of hydrogel beads and barcode beads co-encapsulation. b. Droplets circled by white line were regarded as positive, which means containing only one barcode bead and only one genomic DNA-containing hydrogel bead.

### *Barcode beads quality control and barcode sequences verification*

The barcode beads were synthesized in a DNA synthesizer through a split-pool approach. All barcode beads were quality controlled and verified by probe binding and gradient release before use **(Fig6)**. To verify the sequence on barcode beads, three probes were designed to capture three different regions in the barcode sequence (5’-FAM-GCCGTCTTCTGCTTGGAAAAAAA-3’; 5’-FAM-TCGTATGCCGTCTTCTGC-3’; 5’-FAM-CCGAGCCCACGAGAC-3’), which were hybridized with oligos on beads synthesized through thermal cycling. The barcode beads after hybridization were washed and observed with a fluorescence microscope (**Fig 6a**). To quantify the oligos on barcode beads, ~ 4000 barcode beads were added into a total 40μL reaction system and verified under four different conditions (25℃, 10mM DTT; 25℃, 50mM DTT; 50℃, 10mM DTT; 50℃, 50mM DTT)**.** DTT was used to break the disulfide bond which connected the barcode oligos to polystyrene beads. Temperature also had an impact on releasing the barcode oligos from the polystyrene beads. Higher concentration of DTT and higher temperature could accelerate the releasing speed of oligos from barcode beads (Fig 6b). Hence, the oligos on barcode beads could be completely released during PCR reaction for barcoding each genomic DNA fragment.

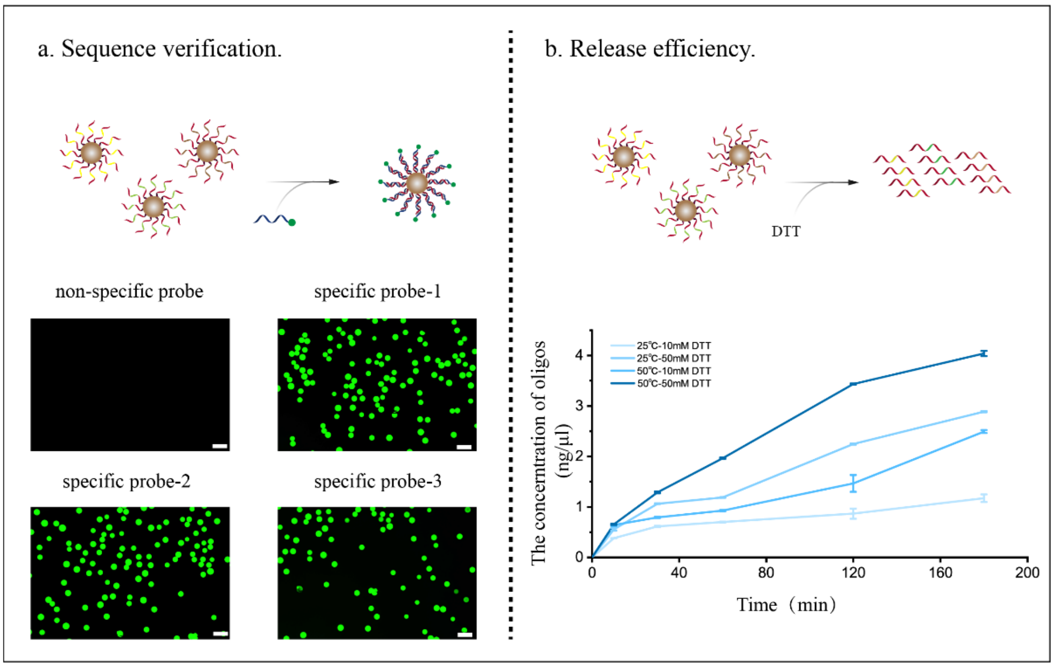


**Fig6: Quality control of barcode beads.** a. Sequence verification of barcode beads with different probes. b. Sequence release efficiency of barcode beads under four conditions.
